# Unique CD8^+^ T Cell Populations Expand during ART and Predict Delayed HIV-1 Rebound

**DOI:** 10.64898/2026.08.11.743993

**Authors:** Jie Wang, Gautam Kundu, Philip K. Ehrenberg, Aviva Geretz, Daina Eiser, Timothy Ezebuiro, Ashan Dayananda, Hiroshi Takata, Carlo Sacdalan, Somchai Sriplienchan, Nittaya Phanuphak, Suteera Pinyakorn, Nelson L. Michael, Lydie Trautmann, Fred Sawe, John W. Mellors, Julie A. Ake, Sandhya Vasan, Rasmi Thomas, the RV254 study team

## Abstract

Antiretroviral therapy (ART) suppresses HIV-1 replication but does not eliminate the latent reservoir, resulting in viral rebound with variable kinetics after treatment interruption. How the immune cell states established during ART influences timing of rebound is not fully understood. In this study, we analyzed 111 participants across multiple cohorts, with 188 single-cell multiomic samples generated and integrated for joint analysis. Longitudinal profiling of peripheral blood mononuclear cells from individuals with acute HIV-1 infection on ART, spanning early infection through sustained therapy and pre-analytical treatment interruption, revealed that time to viral rebound was driven not by global changes in immune composition but by dynamic transcriptional programs within CD8^+^ T cells. During ART, there was a dramatic expansion of a unique cluster of poised naïve CD8^+^ T cells, with a distinct immune state positioned upstream of stem-like memory CD8^+^ T cells along a cell differentiation continuum. The differential abundance of this poised naïve CD8^+^ T cell population was enriched in participants with delayed rebound and showed strong predictive power for discriminating time to rebound. Mechanistically, the poised naïve CD8^+^ T cells exhibited features of a precursor phenotype of stem-like memory CD8^+^ T cells, and showed activation of the TNFα-NF-κB signaling pathway and increased chromatin accessibility at AP-1 motifs. Notably, both poised naïve CD8^+^ T cells and stem-like memory CD8^+^ T cells were consistently enhanced during ART in both acute and chronic infection. In participants who received investigational therapeutic vaccination, the dominant predictive signal shifted downstream along the differentiation trajectory, with stem-like memory CD8^+^ T cells emerging as the primary determinant of delayed rebound. Together, these findings identify a dynamic CD8^+^ T cell state continuum as a central determinant of HIV-1 rebound, even in the absence of antigen-specificity, where ART establishes a predictive poised naïve state that can be further leveraged by vaccination to enhance protective stem-like memory responses.

## Introduction

The advent of antiretroviral therapy (ART) has transformed HIV-1 infection from a fatal disease into a chronic, manageable condition. By effectively suppressing active viral replication, ART reduces plasma viremia to undetectable levels and enables long-term immune recovery^1^. However, ART does not eliminate latently infected cells, allowing the virus to persist in long-lived reservoirs that trigger viremia upon analytic treatment interruption (ATI) and prevents a cure^2,3^. This fundamental limitation has driven intense interest in strategies that would enable people living with HIV-1 (PLWH) to safely discontinue ART. Such approaches broadly aim either to eliminate replication-competent virus (HIV-1 eradication) or to achieve durable immune-mediated control of viremia in the absence of therapy (HIV-1 remission)^4^. Regardless, viral rebound occurs in most individuals within 2-4 weeks following treatment interruption, whether therapy is initiated during acute or chronic HIV-1 infection^5,6^. Treatment interruption studies have demonstrated substantial inter-individual variability in time to viral rebound among PLWH, with a subset termed post-treatment controllers (PTCs) achieving sustained viral suppression in the absence of antiretroviral therapy^7–9^. Prior investigations into determinants of rebound timing have identified HIV-1 reservoir size, timing of ART initiation, pre-ART clinical parameters such as plasma viral load (VL), time to viral suppression (VLS) upon ART initiation, and specific host HLA class I alleles as key modulators^10–15^.

Immune compartments, including T cells, monocytes, natural killer (NK) cells, and the broader innate immune program, have also been linked to reservoir persistence, immune activation, and control of viral replication^16,17^. Our recent studies identified monocytes as central players associated with HIV-1 reservoir size in individuals undergoing ART, and elevated *IL1B* expression in monocytes correlated with smaller reservoir sizes, suggesting a potential immunomodulatory link between innate immune activation and viral rebound^18,19^. NK cells contribute to early antiviral control through cytotoxicity and antibody-dependent cellular cytotoxicity, and their functional competence has been associated with delayed rebound kinetics^17^. CD4^+^ T cell subsets, including T follicular helper cells, serve as key reservoir sites while also orchestrating humoral immunity, linking viral persistence with immune regulation^20,21^. In addition, B cell and antibody responses, particularly broadly neutralizing and Fc-mediated functions, may modulate post-treatment viral control by shaping both viral clearance and immune activation states^22,23^. Evidence for the role of CD8^+^ T cells in HIV-1 control initially emerged from host genetic studies in natural history cohorts^24,25^. Notably, protective HLA alleles previously associated with viral suppression are now also implicated in delayed viral rebound^15^, suggesting that CD8^+^ T cells may also play a critical role in post-treatment control. During acute infection, HIV-specific cytotoxic CD8^+^ T cells exert strong selective pressure on the virus by recognizing and eliminating infected CD4^+^ T cells, contributing to the initial decline in viremia^26,27^. However, cytotoxic CD8^+^ T cells fail to achieve sterilizing immunity, in part due to rapid viral escape mutations and progressive functional exhaustion of the antiviral response^28^. In parallel, studies of HIV-specific CD8^+^ T cells from HIV-1 elite controllers and early treated individuals have highlighted the importance of CD8^+^ T cell stemness beyond cytotoxicity alone^29,30^. These findings support a model in which durable viral control depends not only on immediate effector functions but also on the maintenance of a regenerative CD8^+^ T cell pool. Interrogation of HIV-specific immune responses typically requires substantial cellular input, representing a major limitation in clinical studies incorporating analytical treatment interruption. To overcome this constraint, we sought to define how immune cell states established during suppressive ART shape time to viral rebound at the system level, independent of antigen specificity. In parallel, we aimed to identify robust immunological correlates providing mechanistic insight, as well as are predictive of rebound kinetics that are amenable with low-input sampling and broadly applicable across clinical settings.

Single-cell multiomics technologies now permit comprehensive characterization of the immune landscape with unprecedented resolution^31–33^. This approach is ideal for unbiased analyses and has led to new discoveries^19^. We leveraged the ATI framework within an acute HIV infection (AHI) cohort to investigate the strongest immune determinants of viral rebound across all immune populations in peripheral blood. Longitudinal peripheral blood mononuclear cells (PBMCs) from AHI through ART suppression were analyzed using single-cell Cellular Indexing of Transcriptomes and Epitopes by sequencing (CITE-seq)^34^, and single-cell Assay for Transposase-Accessible Chromatin with Select Antigen Profiling (ASAP-seq)^35^, coupled with plasma proteomics and flow cytometry profiling to discover multi-layered circulating immune states associated with viral rebound. We found that time to viral rebound is not associated with global immune composition change in peripheral blood but is instead associated with transcriptional programs within CD8^+^ T cells. We identified a distinct poised naïve CD8^+^ T cell state that emerges during AHI, and was further expanded under ART, characterized by coordinated transcriptional and chromatin remodeling, including activation of AP-1-and TNFα-NF-κB regulatory programs. This state exhibited features consistent with progenitor-like populations with the potential to differentiate into stem-like memory CD8^+^ T cells, and its abundance was strongly associated with delayed viral rebound. Intervention using therapeutic vaccination shifted the dominant predictive signal toward stem-like memory CD8^+^ T cells, while maintaining the poised features, highlighting a redistribution along the CD8^+^ T cell axis of differentiation continuum under ART, highlighting the potential for harnessing the host immune system for achieving HIV-1 remission.

## Results

### Clinical characteristics and immune remodeling during ART

This study was performed using samples from multiple ATI studies (RV411, RV254, RV405, and RV397) in the RV254 cohort in Thailand. Clinical and demographic characteristics of the study participants are summarized in **Supplementary Table 1.** Briefly, 27 PLWH (range 18 – 44 years) participants received standard ART and were without additional interventions or in the placebo arm of specific interventional studies^6,7,36,37^. Participants were diagnosed during AHI and had been on ART for a median of 2.81 years (range 2.18 – 6.09 years). Longitudinal peripheral blood samples were collected at AHI, 60 weeks post-ART initiation (ART), and immediately prior to ATI (pre-ATI) (**Fig. 1a**). During ATI, all participants experienced viral rebound (viral load (VL) >20 copies/mL), except for one participant carrying HLA-B*57, who did not rebound (last assessed at 183 days post ATI) and was therefore excluded from the downstream analyses.

**Figure 1.**
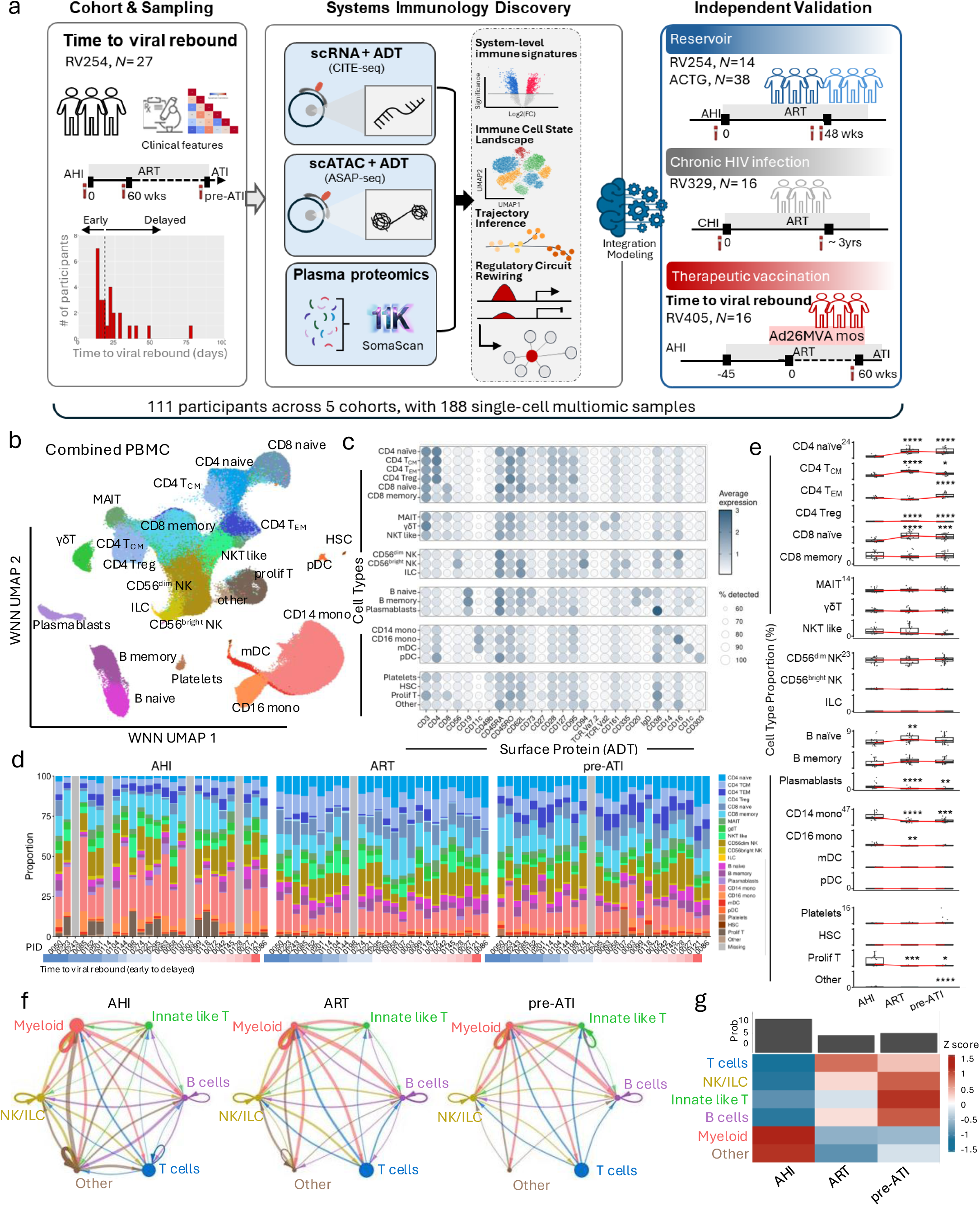
Overview of dynamic changes in immune cells in peripheral blood using multiomics in the context of HIV-1 outcomes. (**a**) Integrated single-cell profiling and prediction modeling was performed on longitudinal peripheral blood mononuclear cells (PBMC) collected across samples from acute HIV infection (AHI), 60 weeks after initiation of antiretroviral therapy (ART), and pre-analytical treatment interruption (pre-ATI) timepoints in the absence of other interventions. Findings were validated in additional cohorts with various HIV-1 outcomes, including reservoir size, chronic HIV infection, and time to rebound after ATI in the context of therapeutic vaccination. (**b**) Integrated PBMC profiled by CITE-seq across AHI, ART, and pre-ATI timepoints. Cells were embedded using Uniform Manifold Approximation and Projection (UMAP) based on weighted nearest neighbor (WNN) analyses, jointly leveraging transcriptomics (RNA) and surface protein (antibody-derived tag, ADT) modalities. Cells are colored by annotated immune cell types. Each point represents an individual cell (n = 412,302). (c) Surface protein expression across immune cell types shown as a dot plot summarizing ADT measurements representing surface expression of markers corresponding to cell types in PBMC. Circle size indicates the fraction of expressing cells and color represents scaled expression levels. (d) Stacked bar plots show the proportion of each cell type per donor at AHI, ART, and pre-ATI timepoints. Participants are ordered by time to viral rebound, from early (blue) to delayed (red). Missing samples from participants PID 0003, 0114, 0128, and 0243 at AHI timepoint; PID 0198 at the ART; and PID 0221 at pre-ATI were excluded due to failure in quality control. (**e**) Plots of indicated immune cell type proportions across individuals and timepoints. Differential proportions were calculated by the Mann–Whitney U-test and significance was FDR corrected. * *P* < 0.05, ** *P* < 0.01, *** *P* < 0.001, and **** *P* < 0.0001. (**f**) Cell-cell communication (CCI) network dynamics across three timepoints. Cell types were grouped as T cells: CD4 naïve, CD4 T_CM_, CD4 T_EM_, CD4 Treg, CD8 naïve, and CD8 memory cells; innate-like T cells: MAIT cells, γδT cells, and NKT like cells; NK/ILC cells: CD56^dim^ NK, CD56^bright^ NK, and ILC; B cells: B naïve, B memory, and Plasmablasts; myeloid cells: CD14 monocytes, CD16 monocytes, mDC, and pDC; others: platelets, HSC, proliferating cells, and other cells. Network graphs depict inferred ligand-receptor interactions among cell populations at AHI, ART, and pre-ATI timepoints. (**g**) Global rewiring of intercellular communication under different conditions. Bars indicate the probability (Prob) of the sum of the signaling strengths of each timepoint. The Heatmap summarizes the signaling pathways of each cell type.

Longitudinal SOMAscan plasma proteomics performed at the AHI and ART timepoints revealed marked enrichment of inflammatory and immune signaling pathways, including interferon-γ response, allograft rejection, and IL-6/JAK/STAT3 signaling in AHI timepoint. In contrast, ART induced a shift toward metabolic and tissue homeostasis programs, including xenobiotic metabolism and fatty acid metabolism pathways (**Supplementary Fig. S1a–b**), consistent with suppression of acute immune activation and establishment of a remodeled immune state.

### Cellular immune state changes and signaling rewiring during ART

To further investigate how cellular immune states are established under ART, we performed longitudinal single-cell CITE-seq on PBMC at three timepoints (**Fig.1a**), yielding a total of 412,302 high-quality cells. Joint integration of transcriptomic and surface protein antibody-derived tag (ADT) data identified 23 immune cell populations (**Fig. 1b-c; Supplementary Fig. S2**), and ART induced a pronounced contraction of inflammatory and differentiated compartments, including the reduction of CD14^+^ and CD16^+^ monocytes and CD4^+^ effector memory T cells (T_EM_). In parallel, the less differentiated populations expanded, with increased frequencies of naïve CD4^+^ T cells, CD4^+^ central memory T cells (T_CM_), and naïve CD8^+^ T cells at both ART and pre-ATI timepoints (**Fig. 1d-e**). This shift reflects a transition from an activated, effector-skewed immune landscape during AHI towards a more quiescent, stemness-and memory-dominated state under ART.

We next examined the rewiring of cell–cell interactions (CCI) across peripheral immune populations using probabilistic modeling to identify significant communications between cell types^38^. We assessed these interactions by collapsing the 23 identified cell populations into five major compartments: myeloid (monocytes and dendritic cells (DC)), T cells (CD4^+^ and CD8^+^ T), B cells (naïve, memory, and plasmablasts), innate-like T cells (NKT-like, γδT, and MAIT), NK/ILC (CD56^bright^ NK, CD56^dim^ NK, and ILC), and other cell populations (platelets, HSC, proliferating cells, and other cells). (**Fig. 1f**). Globally, intercellular signaling was most pronounced during AHI and was substantially attenuated following ART, as observed by the decreased outgoing and incoming signal probabilities across compartments. This reduction was particularly evident in myeloid cells, which exhibited strong signaling activity during AHI that diminished under ART. In parallel, T cell signaling was increased following ART (**Fig. 1f-g**). This shift suggests a transition from myeloid-driven inflammatory signaling during acute infection toward a more T cell-centered communication network under ART, potentially reflecting reorganization of adaptive immune regulation.

### Naïve CD8^+^ T cells exhibit dramatic transcriptomic changes associated with time to viral rebound

Due to the marked global changes in immune signaling during ART, we investigated which immune cell populations had the strongest association with time to viral rebound upon ATI. The twenty-six participants were classified as early or delayed rebounders based on the median time to viral rebound (19 days; range 13 – 77 days) (**Fig. 2a**). The comparative analyses between early and delayed rebounders revealed broadly similar cell type proportions at the three timepoints (**Supplementary Fig. S3**), suggesting that bulk compositional differences alone do not account for rebound heterogeneity. We therefore hypothesized that time to viral rebound is governed by transcriptional rewiring within cellular states rather than by shifts in population frequencies. To identify clinical variables associated with time to viral rebound in the absence of therapeutic interventions, we applied a SHAP (SHapley Additive exPlanations)^39^-based support vector regression (SVR) model, which identified time to viral suppression as the strongest negative predictor of time to rebound in the absence of other intervention (**Fig. 2b**). We therefore incorporated time to viral load suppression (VLS) as a covariate, together with participant identity, while modeling time to rebound as a continuous variable^40^.

**Figure 2.**
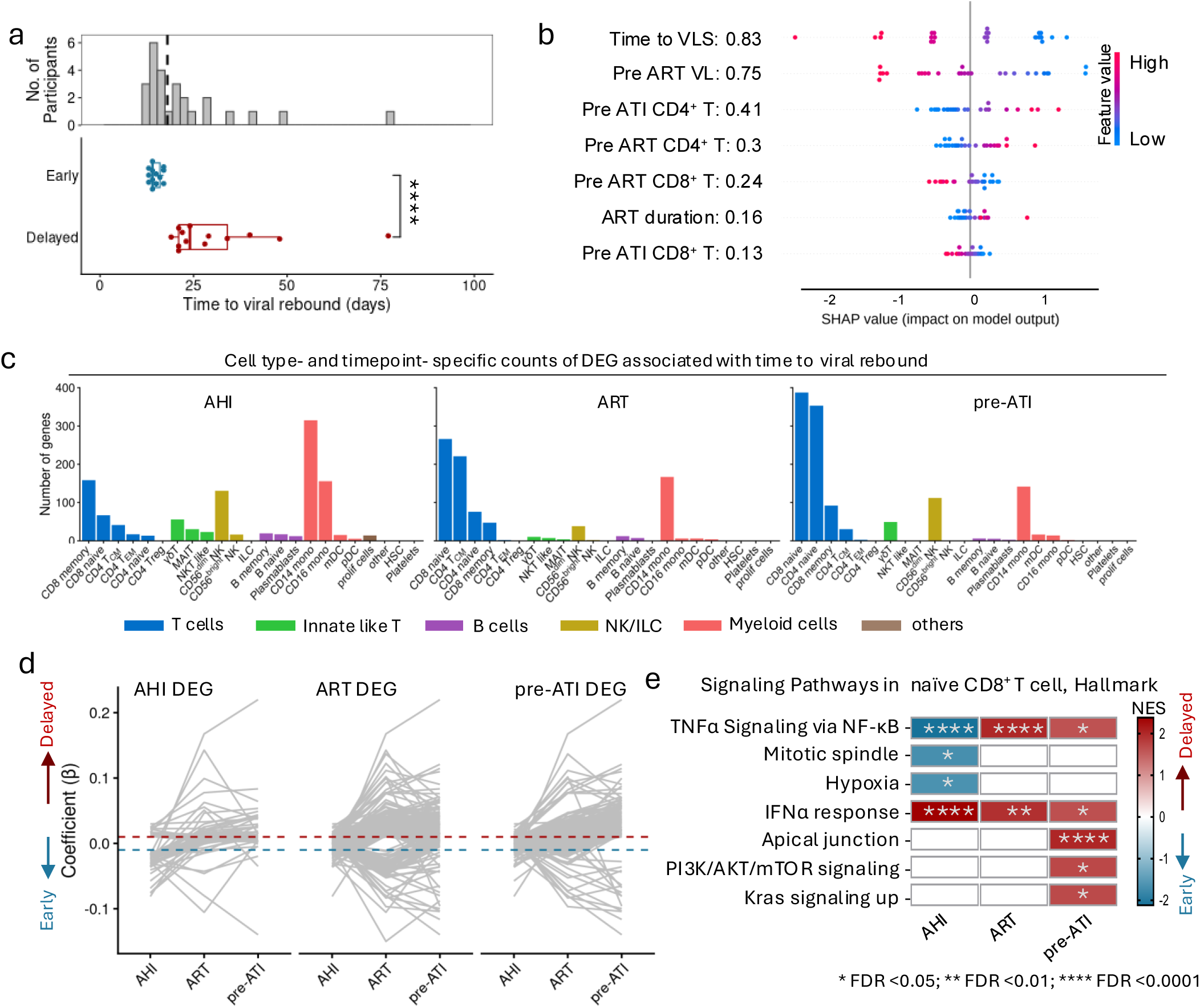
Clinical features and transcriptomic changes associated with time to viral rebound. **(a)** Bar graph (top) shows distribution of time to viral rebound (days) after ATI in RV254 (N=26). Box plot (bottom) shows the participants classified into delayed or early rebound categories based on the median value of 19 days. Statistical significance was determined by a two-sided Mann–Whitney U-test. **** *P* < 0.0001. **(b)** An SVM (Support Vector Machine) model was trained using clinical features to predict time to viral rebound following ATI. Features are ranked along the y-axis in descending order of their global importance (mean absolute SHAP value). Each point represents an individual participant. Positive SHAP values indicate a contribution to delayed time to viral rebound, while negative SHAP values indicate a contribution to early time to viral rebound. The color of each point represents the actual value of that feature. Pink indicates high, while blue indicates low values. VLS: Viral Load Suppression; VL: Viral Load. **(c)** Bar plots depict the number of differentially expressed genes (DEG) associated with time to viral rebound per cell type at AHI, ART, and pre-ATI. Differential expression analyses were performed using MAST, adjusting for participant ID (PID) and time to VLS as covariates. Genes meeting the significance thresholds of |β| > 0.01 and Bonferroni-adjusted *P* < 0.05 were included. Complete DEG lists for each timepoint are provided in Source Data (Extended Table). **(d)** Dynamic associations of significant DEG in naïve CD8^+^ T cells with time to viral rebound at each timepoint. Positive and negative values indicate the association with either delayed or early rebound, respectively. **(e)** The heatmap displays pathway enrichment analyses in naïve CD8^+^ T cells at three timepoints. Color represents normalized enrichment scores (NES). All pathways shown are significantly enriched (FDR adjusted *P* <0.05) and are associated with the either delayed (red) or early (blue) time to viral rebound. * *P* < 0.05, ** *P* < 0.01, **** *P* < 0.0001.

At AHI, the strongest rebound-associated transcriptional signals were observed in myeloid cells, particularly CD14^+^ monocytes, which exhibited the highest number of differentially expressed genes (DEG) (**Fig. 2c**, **Supplementary Fig. S4**). While previously described by us and others, this timepoint is largely confounded by elevated gene expression of interferon stimulated genes, cytokines and IFN-resistant phenotypes due to changes in viremia during acute infection^18,41–44^ (**Extended Table**). By contrast, when viremia was suppressed during ART, we observed that the rebound-associated transcriptional variation shifted towards lymphoid components. Notably, naïve CD8^+^ T cells emerged as the dominant population, harboring the largest number of DEG at both ART (266) and pre-ATI (387) timepoints (**Fig. 2c**, **Supplementary Fig. S4** and **Extended Table**). DEG identified in naïve CD8^+^ T cells at the AHI timepoint showed either reversed or no association with time to viral rebound during ART, while DEG that were identified at ART and pre-ATI were largely concordant and predominantly upregulated in delayed rebounders (**Fig. 2d**, **Extended Table**).

Naïve CD8^+^ T cells have traditionally been considered quiescent, yet recent single-cell studies reveal substantial heterogeneity in healthy adults^45^. Consistent with this, gene set enrichment analyses (GSEA)^46^ revealed dynamic, time-dependent pathway rewiring. Interferon-α response signatures were associated with delayed rebound at AHI but were attenuated, though still significant, during ART (**Fig. 2e**; **Supplementary Fig. S5a-b**). In contrast, TNFα -NF-κB signaling was enriched in early rebounders at AHI but shifted toward delayed rebounders during ART, persisting through the pre-ATI timepoint. Additionally, pathways related to apical junction and PI3K/AKT/mTOR signaling only emerged at the later stage of ART (pre-ATI). Collectively, these findings define a temporal reprogramming of naïve CD8^+^ T cell states, transitioning from an early interferon-driven inflammatory program to a more metabolically-and signaling-poised state during prolonged ART.

### ART reveals a continuous spectrum of CD8^+^ T cell differentiation states

Given the pronounced transcriptomic rewiring of naïve CD8^+^ T cells associated with time to viral rebound, we next examined how ART reshapes CD8^+^ T cell phenotypes at the population level by comparing population frequencies from flow cytometry at the AHI and pre-ATI timepoint. During ART, there were significant increase in the proportions of naïve (TN, CD45RA^+^CD27^+^CCR7^+^) and central memory (T_CM_, CD45RA^-^CD27^+^CCR7^+^) populations, accompanied by a reduction in transitional memory (T_TM_, CD45RA^-^CD27^-^CCR7^+^) cells, while effector memory (T_EM_, CD45RA^-^CD27^-^CCR7^-^) and terminally differentiated (T_TD_, CD45RA^+^CD27^-^CCR7^-^) populations remained largely unchanged (**Fig. 3a-b; Supplementary Fig. S6a**). Despite this global shift, canonical flow cytometry gating did not reveal significant differences in CD8^+^ T cell subset frequencies between early rebounders and delayed rebounders at the pre-ATI timepoint (**Supplementary Fig. S6b)**.

**Figure 3.**
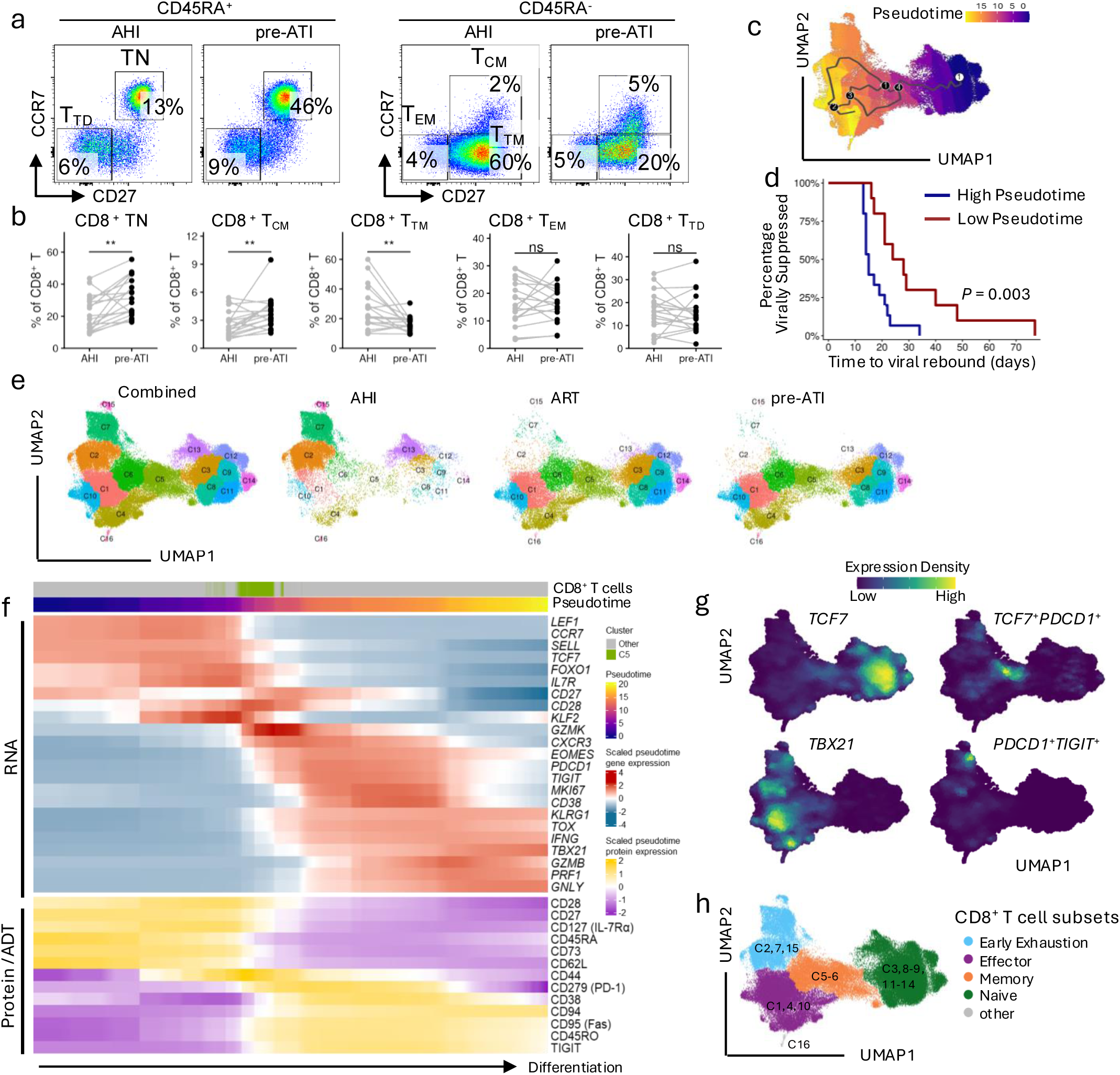
CD8^+^ T cell pseudotime trajectory reveals ART-induced differentiation impacting time to viral rebound. **(a)** Flow cytometry gating strategy for CD8^+^ T cell subsets at the AHI and ART timepoints from a subset of the RV254 ATI study (N=18). CD8^+^ T cells were classified into naïve (CD8^+^ TN), terminally differentiated (CD8^+^ T_TD_), effector memory (CD8^+^ T_EM_), transient memory (CD8^+^ T_TM_), and central memory (CD8^+^ T_CM_) populations based on cell surface markers. **(b)** Paired comparisons of the frequencies of CD8^+^ T cell subsets defined in (a). Each dot represents an individual participant, with lines connecting paired samples at AHI (gray) and post-ART (black). Statistical significance was determined by Wilcoxon matched-pairs signed rank test. ** *P* < 0.01, ns: not significant. **(c)** UMAP of CD8^+^ T cells constructed from the CITE-seq RNA assay and colored by pseudotime as inferred using Monocle 3. Pseudotime ordering reflects a continuum of cellular differentiation along an inferred trajectory with lower values indicating less differentiated states. **(d)** Survival analysis of the association of CD8^+^ T pesudotime and time to viral rebound at the pre-ATI timepoint. **(e)** UMAP of CD8^+^ T cell subclusters (C1-C16) along the inferred pseudotime trajectory, split by time points. **(f)** Pseudotime expression heatmap representing CD8^+^ T cell differentiation. The upper section of the heatmap illustrates the density distribution of expression of key CD8^+^ T cell RNA along the pseudotemporal trajectory. The lower section of the heatmap illustrates the density distribution of expression of key CD8^+^ T cell surface markers along the pseudotemporal trajectory. Cells are ordered by pseudotime; T_SL-M_ C5 cells are highlighted in green. Expression levels are scaled within each marker. (**g**) Density plots show the expression of key transcriptomic features of CD8^+^ T cells along the trajectory. (**h**) UMAP showing Monocle-inferred CD8^+^ T cell trajectory, with distinct cell subsets colored as indicated.

Further subclustering of naïve CD8^+^ T cells using hierarchical clustering ^47^ did not identify distinct subpopulations associated with rebound. Therefore, in order to capture finer-grained cellular states, we performed pseudotime analyses of single-cell naïve and memory CD8^+^ T cells across all three timepoints using Monocle3^48–50^. The inferred trajectory recapitulated differentiation from naïve toward progressively differentiated states (**Fig. 3c)**. Notably, individuals whose CD8^+^ T cells exhibited more advanced differentiation overall (higher pseudotime values) tended to experience earlier viral rebound (*P* = 0.003) (**Fig. 3d**).

Unsupervised clustering along this continuum identified 16 transcriptionally distinct subclusters (**Fig. 3e**). Integrated transcriptomic and surface protein annotations revealed a progressive transition from naïve CD8^+^ T cells to effector memory states. At the RNA level, canonical naïve markers, including *SELL*, *LEF1*, *CCR7*, and *TCF7* (TCF-1), were highly expressed early in the trajectory and gradually declined as cells differentiated (**Fig. 3f**). In contrast, effector-associated genes such as *TBX21* (T-bet), *IFNG*, *PRF1* (Perforin), and *GNLY* showed progressive increases along the change in pseudotime. Notably, cluster 5 (C5) appeared to represent a bridge sharing both naïve and effector signatures (**Fig. 3f**). Consistent patterns were also observed at the protein level. Naïve surface markers, including CD45RA and CD62L, were highly expressed early in the trajectory and diminished with differentiation, whereas effector and memory-associated markers such as CD45RO, CD94, and CD95 (Fas) were progressively enriched toward the terminal states. C5 again displayed shared naïve and effector features, characterized by high CD44 expression (**Fig. 3f**), consistent with a stem-like memory T cell (T_SL-M_) phenotype. Based on these molecular features, along with co-expression patterns observed in the feature plots (**Fig. 3g**), clusters were further grouped into major functional categories: CD8^+^ naïve T cells (TN; C3, C8, C9, C11–C14: *TCF7^+^),* stem-like memory T cells (T_SL-M_; C5: *TCF7^+^PDCD1^+^*), central memory T cells (T_CM_; C6), early exhausted T cells (C2, C7, C15: *PDCD1^+^TIGIT^+^),* effector memory and terminally differentiated T cells (T_EM_ and T_TD_; C1, C4, C10: *TBX21^+^*), and a minor outlier population (C16) (**Fig. 3g-h; Supplementary Fig. S7**).

### ART-induced poised naïve-like cluster drives dynamic remodeling within the CD8^+^ T cell population

Since CD8^+^ T cell differentiation status associated with time to viral rebound, we wanted to understand which specific subcluster(s) contributed to this outcome. Interestingly, individuals with delayed rebound exhibited a significantly higher frequency of C3 at pre-ATI (*P* < 0.05), whereas C6 displayed the opposite trend (*P* < 0.05) (**Fig. 4a)**. Furthermore, the regression analysis demonstrated that higher frequencies of C3 were strongly associated with increased time to viral rebound (*FDR* = 0.003), while the effect of C6 was not significant (**Fig. 4b**). Therefore, we focused subsequent analyses on C3. Although CD8^+^ TN C3 retains canonical naïve features, including expression of *TCF7* and *LEF1* at the transcriptomic level and CD45RA^+^ and CD45RO^-^surface protein phenotypes, its transcriptomic profile closely aligns with T_SL-M_ cells (C5) (*rho* = 0.72, *P* < 0.05), suggestive of a precursor memory-poised state (**Fig. 4c**). Partition-based graph abstraction (PAGA)^51^ further supported this relationship, as CD8^+^ TN C3 showed a strong connectivity to T_SL-M_ (C5), consistent with a memory-oriented trajectory (**Fig. 4d**).

**Figure 4.**
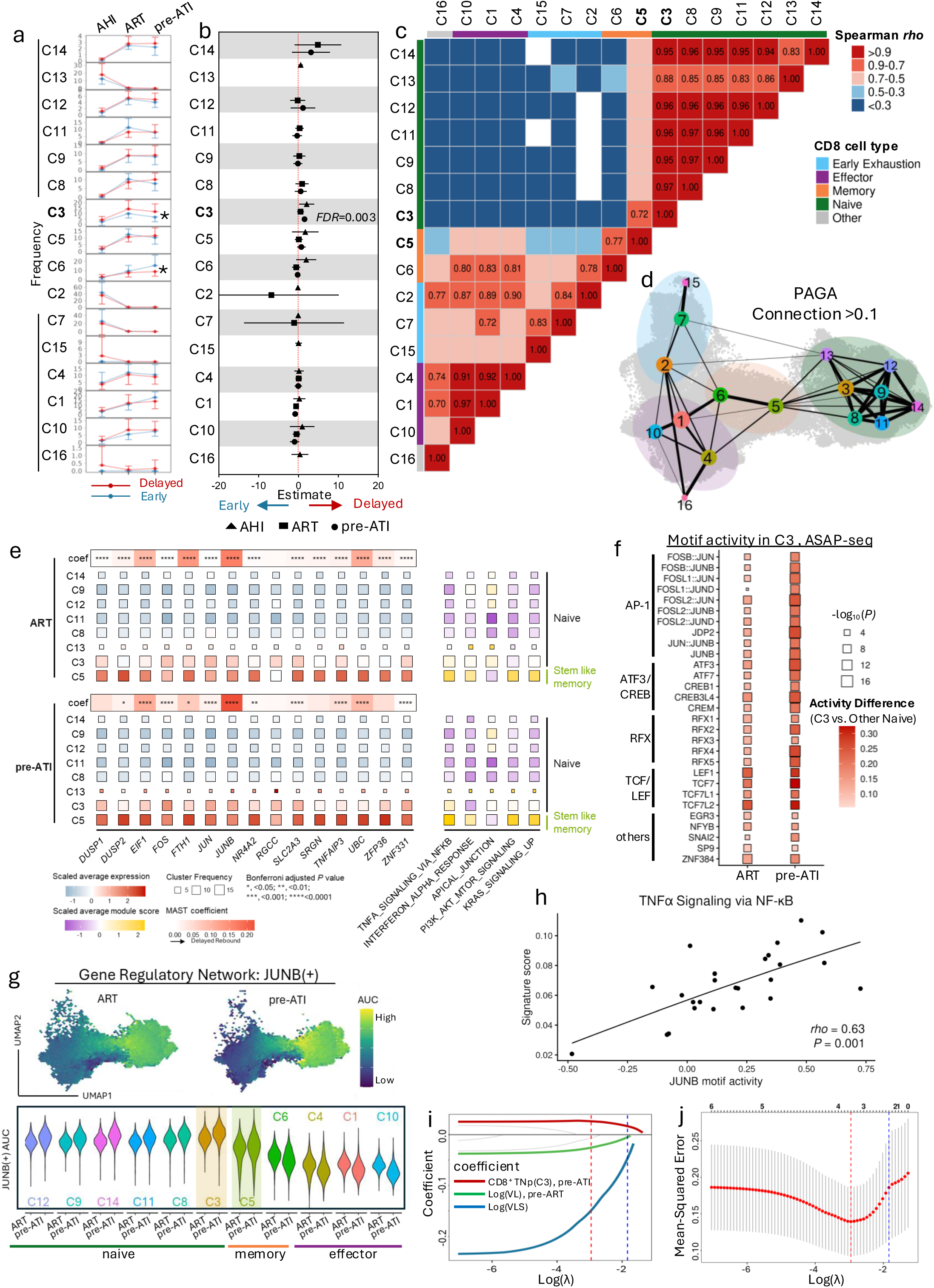
A subset of naïve CD8^+^ T cell predicts time to viral rebound. **(a)** The line plots summarizing the comparative CD8^+^ T cell subcluster cell proportions between early (blue) vs delayed (red) rebound at three time points. Statistical significance was determined by a two-sided Mann–Whitney U-test. * *P* < 0.05. **(b)** Univariate regression analyses showing the estimated effect of CD8^+^ T cell subclusters on time to viral rebound at three timepoints. Subclusters C14 at AHI; and C13, C15, and C16 at both ART and pre-ATI timepoints were excluded due to low frequencies (<1% of CD8^+^ T cells); triangles, squares, and circles represent the AHI, ART, and pre-ATI timepoints, respectively. *P* values were adjusted for multiple testing using FDR across all subclusters and timepoints. **(c)** The heatmap shows the Spearman correlations between CD8^+^ T cells subclusters with the top 20 DEG extracted from each cluster (sorted by fold change of gene expression with adj *P* < 0.05); red indicates high correlation, and blue indicates low correlation. Only correlations with *rho* > 0.7 and *P* < 0.05 have values indicated. **(d)** Partition-based Graph Abstraction (PAGA) analysis graph depicts inferred lineage relationships among clusters; edge width reflects connectivity strength. **(e)** Differential gene expression between C3 and all other naïve clusters (≥200 cells) was assessed. Genes with FDR adjusted *P* < 0.05 and logFC > 0.05 in C3 versus every other naive cluster at both treatment timepoints were selected. Heatmaps (left panel) display log-normalized expression, with color indicating z-scaled average expression across cluster (other CD8^+^ TN, C3, and C5)–timepoint combinations; color (blue-white-red) represents the scaled gene average expression. Heatmaps with MAST coefficient (coef) for each feature with respect to time to viral rebound; color (white-red) denotes associations with delayed rebound. Bonferroni adjusted *P* value: * *P* < 0.05, ** *P* < 0.01, *** *P* < 0.001, **** *P* < 0.0001. Heatmap (right panel) displays the pathway module scores calculated across CD8^+^ TN cell subsets (C3, C8, C9, C11, C12, C13, and C14) and C5 at three time points. Size of the square reflects the proportion of cells per cluster, and color (purple/white/yellow) represents the scaled module score. **(f)** Heatmap of top enriched transcription factors in C3 vs. other CD8^+^ TN at ART and pre-ATI time points based on ASAP-seq chromatin accessibility analyses. Transcription factors are grouped into AP-1, ATF/CREB, RFX and TCF/LEF families. Box size indicates −log₁₀ (*P* value), and color represents average differential activity between C3 vs. other naïve CD8^+^ T cells. **(g)** UMAPs show JUNB regulon activity (AUC scores) in CD8^+^ T cells at ART and pre-ATI timepoints (top). Violin plots (bottom) show the distribution of JUNB AUC scores across CD8^+^ T cell subclusters (restricted to clusters with >50 cells at both timepoints). (**h**) Spearman correlations between pathway signature scores of TNFα Signaling via NF-κB and JUNB activity. **(i)** Trajectories of LASSO coefficients for five clinical features as a function of the regularization parameter (log(λ)). Vertical dashed lines indicate selected λ values. Clinical features including CD4^+^ T cell counts, CD8^+^ T cell counts, and ART duration that shrank to zero before λ_min were grayed out. **(j)** Cross-validation for model selection. Mean squared error across λ values, with vertical lines indicating the minimum error and the 1-SE rule for model parsimony.

Based on the elevated abundance of the CD8^+^ TN C3 subset in participants with delayed rebound at pre-ATI (**Fig. 4a-b**), we then defined its functional properties via differential gene expression analyses comparing C3 vs. other individual CD8^+^ TN clusters (C8, C9, C11, C12, C13, C14). At the transcriptomic level, CD8^+^ TN C3 was distinguished by elevated expression of immediate and early activation-responsive gene families, such as *FOS*, *JUN*, *JUNB*, *NR4A2*, *ZFP36*, *TNFAIP3*, as well as the metabolic priming genes *SLC2A3, EIF1,* and *FTH1,* especially at pre-ATI (**Fig. 4e**). *JUNB*, *JUN*, *FOS*, *DUSP2*, *EIF1*, *SLC2A3*, *TNFAIP3*, and *UBC* expression also showed significant associations with delayed rebound in the total naïve CD8^+^ T cell population (Bonferroni adjusted *P* value < 0.05; MAST coefficient > 0.01) (**Fig. 4e; Extended Table**). Notably, these activation-responsive features were further enhanced as cells transitioned into the stem-like memory state (T_SL-M_, C5). Moreover, when comparing with other naïve subclusters, (**Fig. 2e**), C3 exhibited significant enrichment in TNF-α signaling via NF-κB, PI3K-AKT-mTOR signaling, and Kras signaling pathways during ART, and those pathways were also enhanced in C5 (**Fig. 4e**). This activation-responsive yet naïve-like signature, coupled with positioning along a memory-like trajectory, suggests C3 is a poised naïve-like CD8^+^ T cell subset, hereafter referred to as CD8^+^ TNp, characterized by heightened activation readiness that is further amplified upon transition to the T_SL_-_M_ state, and a novel contributor to delayed time to viral rebound.

### CD8^+^ TNp undergo progressive epigenetic and regulatory priming via AP-1-centered transcriptional networks

To investigate mechanisms underlying CD8^+^ TNp epigenetic regulation, we performed ASAP-seq, enabling simultaneous profiling of chromatin accessibility and surface protein expression^35^. Broad PBMC cell types (total cells = 305,487) were annotated (**Supplementary Fig. S8**), and subsequent analyses were focused on CD8^+^ T cells across all three timepoints. Pseudotime values and cluster annotations from CITE-seq (**Fig. 3c, e**) were mapped onto the ASAP-seq dataset, and dynamic shifts in transcription factor accessibility (in genes *TCF7* and *TBX21*) supported the previously identified pseudotime structure (**Supplementary Fig. S9a-b**). Corroborating the transcriptional analysis, higher CD8^+^T cell pseudotime at pre-ATI was associated with earlier viral rebound (*P* = 0.039), while increased abundance of CD8^+^ TNp was strongly associated with delayed rebound (*P* = 0.008) (**Supplementary Fig. S9c-d**).

Focusing on the CD8^+^ TNp, we observed selective enrichment of chromatin accessibility at AP-1, ATF3/CREB, TCF/LEF, and RFX family motifs, compared with other naïve clusters, with AP-1 being predominant (**Fig. 4f**). Most of these signals, including AP-1, were stronger at the pre-ATI timepoint, suggesting progressive epigenetic priming. Multiomic integration showed that the AP-1 family motif accessibility was positively correlated with the previously calculated gene module scores of delayed rebound in naïve CD8+ T cells, at both ART (*rho* = 0.55, *P* = 0.005) and pre-ATI (*rho* = 0.49, *P* = 0.01) timepoints (**Supplementary Fig. S9e-f**). These findings implicate AP-1 as a key driver of naïve CD8^+^ T cells remodeling linked to delayed time to viral rebound. Consistently, gene regulatory network (GRN) analysis revealed JUNB regulon activity in CD8^+^ TNp, especially at pre-ATI, with a transition through T_SL-M_, followed by marked decline upon differentiation into memory/effector states (**Fig. 4g**). We further found that JUNB motif activity in CD8^+^ TNp at the pre-ATI timepoint was positively associated with TNFα-NF-κB signaling pathway (*rho* = 0.63, *P* =0.001) (**Fig. 4h**).

Collectively, these results support a model in which CD8^+^ TNp undergoes stepwise epigenetic priming driven by AP-1-centered regulatory networks during ART, with progressively increasing accessibility at key inflammatory and activation-associated motifs. Importantly, the convergence of AP-1-dominated regulatory programs in both C3 and C5 subsets suggests that this priming process represents an early transcriptional and epigenetic continuum that may contribute to the establishment of downstream stem-like memory CD8^+^ T cell states during ART.

### CD8^+^ TNp cells predict time to viral rebound

To assess the clinical relevance of CD8^+^ TNp, we integrated immunological and clinical features, using least absolute shrinkage and selection operator (LASSO)-penalized Cox regression^52^. As the regularization parameter (λ) increased, coefficients for non-informative features shrank to zero (λ_min = 0.05, λ_1se = 0.16; **Fig. 4i-j**), enabling selection of the most predictive features. Cross-validation identified three features at the optimal λ_min – time to VLS, pre-ART VL, and pre-ATI CD8^+^ TNp frequency – that collectively minimized partial likelihood deviance (**Fig. 4i-j**), while counts of CD4^+^ and CD8^+^ T cells, as well as ART duration, did not contribute to prediction of time to viral rebound (**Fig. 4i-j**). Moreover, pre-ART VL and time to VLS contributed negative associations to the model, while pre-ATI CD8^+^ TNp frequency contributed to a positive association. Further, removing CD8^+^ TNp from the model markedly impaired predictive performance (**Supplementary Fig. S10a-b**), confirming its dominant contribution and highlighting it as a critical determinant of time to viral rebound. These findings establish that pre-ATI CD8^+^ TNp frequency robustly predicts delayed viral rebound.

### Validation of unique T cell populations in independent HIV-1 studies and outcomes

To extend and validate our findings, we evaluated the transferability of the identified CD8^+^ TNp cell functional program to other independent orthogonal studies of HIV-1 phenotypes such as reservoir size, which is a key determinant of viral rebound following ATI^12,53^. We transferred our CD8^+^ T cell annotations onto two ART cohort scRNA-seq datasets having total HIV DNA measurements, an independent subset of RV254 participants from Thailand (N=14) and the A5354 cohort from the Americas (N=38)^18^. All participants initiated ART during Fiebig stages III-V with samples from both studies having total HIV DNA measurements at week 48 after ART initiation (**Fig. 5a**). While total HIV DNA includes both intact and defective proviruses and thus represents an overestimate of the replication-competent reservoir, prior studies have shown that total and intact HIV DNA are strongly correlated^54^. For the RV254 cohort, we also had scRNA-seq and clinical parameters from the AHI timepoint. We observed that both CD8^+^ TNp C3 and its downstream subset, CD8^+^ T_SL-M_ C5, are significantly expanded in RV254 after 48 weeks of ART (**Fig. 5b-c**). Notably, the degree of CD8^+^ TNp expansion from AHI to the ART timepoint was strongly associated with the rate of HIV DNA decay (*rho* = 0.82, *P* = 0.0004) (**Fig. 5d**). In addition, CD8^+^ TNp frequencies at the ART timepoint showed a modest increase in individuals with undetectable HIV DNA reservoirs, although this trend did not reach statistical significance (**Supplementary Fig S11**). However, in A5354, a larger independent cohort, higher frequencies of CD8^+^ TNp C3 were significantly associated with smaller reservoir size (*rho* = −0.36, *P* = 0.026) (**Fig. 5e-f, Supplementary Fig S11**). Together, these results support a robust relationship between ART-induced CD8^+^ TNp expansion and reduced HIV-1 reservoir burden.

**Figure 5.**
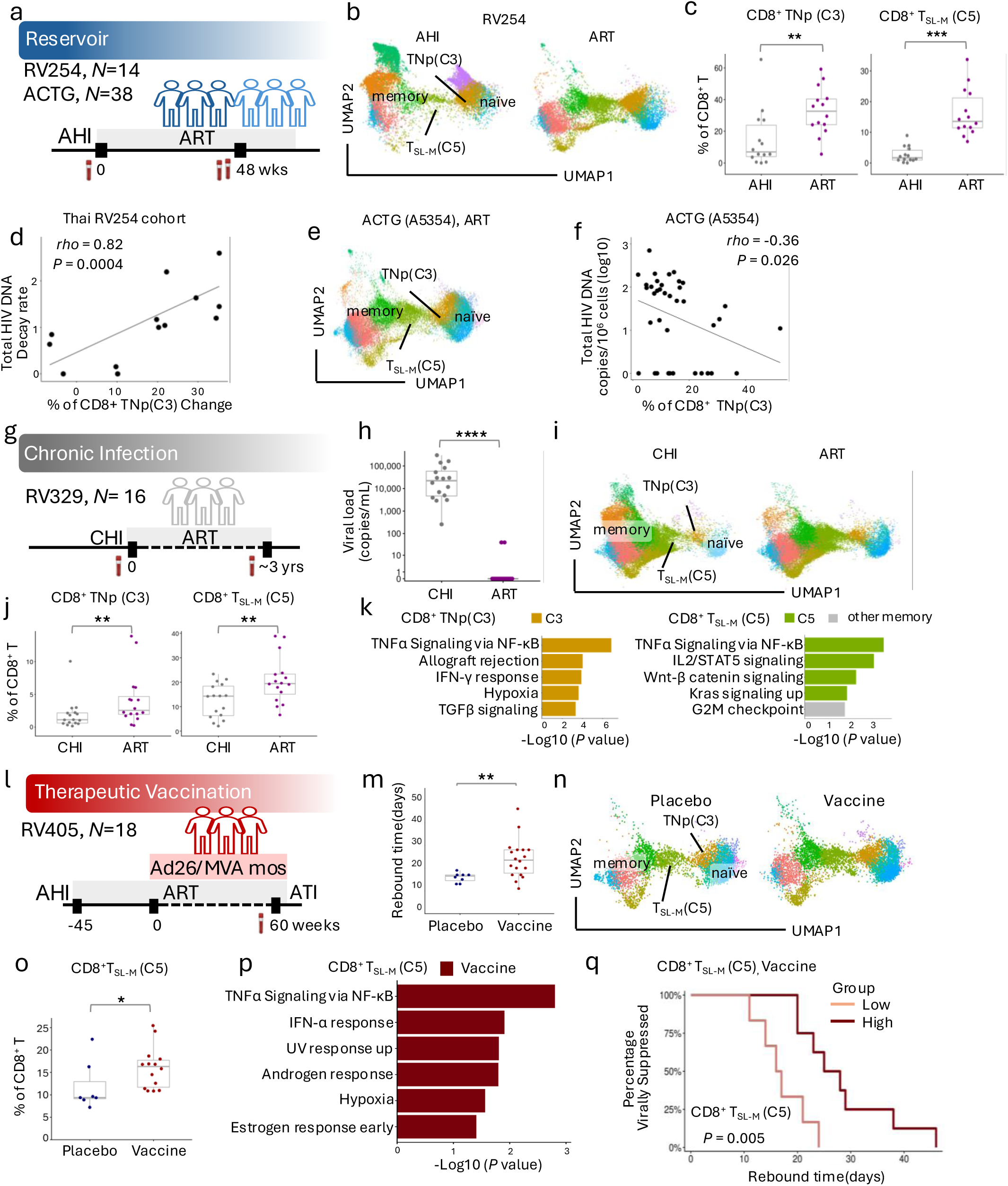
Validation of unique CD8^+^ T cell clusters in multiple independent cohorts. **(a)** Study design of RV254 (N=14) and ACTG A5354 (N=38) with reservoir outcomes. **(b**) RV254 cohort CD8^+^ T cell trajectory mapping at both AHI and ART timepoints. UMAP transferred from Fig. 3c and e to map subclusters. **(c)** Frequencies of CD8^+^ TNp (C3) and CD8^+^ T_SL-M_ (C5) cells were compared in participants from RV254 before and after 48 weeks on ART. Statistical significance was determined by the Wilcoxon matched-pairs signed rank test. ** *P* < 0.01, *** *P* < 0.001. **(d)** Correlation of CD8^+^ TNp (C3) expansion (change in frequency from AHI to ART) and HIV DNA decay rate (log_10_ (Total HIV DNA +1)) from the AHI to ART timepoint in RV254 participants. (**e**) CD8^+^ T cell trajectory mapping at the week 48 time point after ART initiation in the ACTG study. (**f**) Correlation of CD8^+^ TNp (C3) frequency associated with HIV DNA at the ART timepoint in ACTG. **(g)** Study design for RV329 chronic infection (CHI) participants from Kenya (N=16). **(h)** Plasma viral load (copies/ml) in individuals with CHI before ART initiation and after ART. Statistical significance was determined by the Wilcoxon matched-pairs signed rank test. **** *P*<0.0001. **(i)** CD8^+^ T cell trajectory mapping at CHI and ART timepoints. **(j)** Frequencies of CD8^+^ TNp (C3) and CD8^+^ T_SL-M_ (C5) cells were compared in individuals with CHI before ART initiation and during ART. Statistical significance was determined by the Wilcoxon matched-pairs signed rank test. ** *P* < 0.01. **(k)** Bar plots showing representative significantly enriched pathways in CD8^+^ TNp C3 cells, when compared with other CD8^+^ TN (left); and representative significantly enriched pathways in CD8^+^ T_SL-M_ C5 cells, when compared with other memory CD8^+^ T (right); **(l)** RV405 therapeutic vaccine study in participants from Thailand. **(m)** Box plot shows the rebound time comparison between participants receiving the Ad26/MVA mosaic (mos) vaccine (N=18) or placebo (N=8). Statistical significance was determined by the Mann–Whitney U-test. ** *P*<0.01. **(n)** CD8^+^ T cell subcluster mapping. UMAPs of predicted CD8^+^ T cell clusters in the placebo and vaccine groups (prediction confidence > 0.4). Samples from participants PID 0212, 0119, 0304, and 0257 from the vaccination group; and 0221 from the placebo group were removed due to missing sequencing data or failed quality control. (**o**) Box plots showing the relative abundance of CD8^+^ T_SL-M_ C5 cells in placebo versus Ad26/MVA mosaic vaccine recipients. Statistical significance was determined by the Wilcoxon matched-pairs signed rank test. * *P* < 0.05. **(p)** Bar plots showing representative significantly enriched pathways in CD8^+^ T_SL-M_ C5 cells in vaccine versus placebo groups. **(q)** Survival analyss of the association of enrichment of CD8^+^ T_SL-M_ C5 cells in PBMC and time to viral rebound in the Ad26/MVA mosaic vaccine arm (N=14).

Further, to determine whether similar CD8^+^ T cell programs are also engaged during chronic HIV-1 infection (CHI), we analyzed samples from an independent cohort from Kenya, Africa (RV329)^55^, comprising individuals diagnosed during CHI with matched samples after ART initiation (ART) using CITE-seq (**Fig. 5g-h**). Projection of the CD8^+^ T cell differentiation framework from the AHI onto the CHI dataset revealed a marked expansion of both the CD8^+^ TNp C3 and its downstream subset-CD8^+^ T_SL-M_ C5 following ART (**Fig. 5i-j**). Consistent with the observation in the AHI cohort, CD8^+^ TNp cells exhibited significant enrichment of TNFα signaling via the NF-κB pathway relative to the rest of the naïve CD8^+^ T cells, as did CD8^+^ T_SL-M_ cells relative to the rest of the memory CD8^+^ T cells (**Fig. 5k**), demonstrating that ART drives a shared CD8^+^ TNp-to-T_SL-M_ cell trajectory across both stages of acute and chronic HIV-1 infection. Notably, both populations share transcriptional features consistent with poised, activation-ready, and stress-adaptive states, suggesting a common functional program underlying ART mediated CD8^+^ T cell remodeling.

### Therapeutic vaccination increased the frequency of CD8^+^ T_SL-M_ and delayed rebound

To determine if a therapeutic vaccination could elicit these protective CD8^+^ T cell programs during ART, we focused our attention on individuals treated during AHI and maintained on long-term ART who also received an Ad26/MVA mosaic vaccine (**Fig. 5l**). Consistent with prior reports^7^, vaccinated participants exhibited delayed viral rebound following ATI compared to placebo-treated individuals (**Fig. 5m**). CD8^+^ T cell trajectory mapping showed that both CD8^+^ TNp and CD8^+^ T_SL-M_ were present in the placebo and intervention arms. There were no differences in frequencies of CD8^+^ TNp, but the CD8^+^ T cells from the vaccine group exhibited a marked expansion of CD8^+^ T_SL-M_ (C5) compared with placebo **(Fig. 5n-o**). Functionally, CD8^+^ T_SL-M_ cells from vaccinated individuals exhibited pathway enrichment of TNFα signaling via NF-κB, hypoxia, UV response up, and interferon-α response pathways (**Fig. 5p**), consistent with a functionally poised and metabolically adaptable state. More importantly, within the vaccine group we observed that the abundance of CD8^+^ T_SL-M_ cells was significantly associated with delayed rebound (*P* = 0.005) (**Fig. 5q**), suggesting that the T_SL-M_ population plays a key role in mediating post-treatment viral control and can be elicited in the setting of vaccination. This association was stronger than that of the other clinical parameters such as pre-ART viral load (*P* = 0.15) and time to VLS (*P* = 0.04) (**Supplementary Fig. S12a-b**). Collectively, these findings suggest that therapeutic vaccination reshapes the cellular correlates of delayed rebound, shifting the dominant association from CD8^+^ TNp cells during ART specifically towards CD8^+^ T_SL-M_ cells.

## DISCUSSION

HIV-1 infection profoundly perturbs the immune system both phenotypically and functionally. Although ART effectively suppresses viral replication and prevents disease progression, residual immune activation persists, and lifelong treatment remains necessary due to the persistence of viral reservoirs^56^. Understanding the state of immune readiness prior to ART interruption is therefore critical for advancing HIV-1 cure strategies, particularly those aimed at achieving durable remission without continuous therapy. In this context, defining the immune features that distinguish individuals who maintain viral control from those who rapidly rebound, remains a central challenge. Here, we applied a longitudinal, paired, and integrated multiomics framework to comprehensively characterize circulating immune states across diverse infection and treatment contexts. Our results demonstrate that ART induced a coordinated remodeling of the immune system, transitioning the systemic environment from an inflammatory to a metabolically oriented state, while shifting cellular composition toward less-differentiated immune subsets. Notably, this remodeling prominently affected the naïve CD8^+^ T cell compartment, which emerged as a key determinant of time to viral rebound after ART interruption. Specifically, we identified a poised naïve-like CD8^+^ T cell subset characterized by activation-associated AP-1 and NR4A transcriptional programs, stress-adaptive signaling, and partial memory/cytotoxic priming, which was enriched during ART and associated with delayed viral rebound.

Our prior work demonstrated that CD14^+^ monocytes exhibit robust interferon-driven responses associated with a smaller reservoir size at 48 weeks of ART, which in turn is closely linked to time to rebound^18^. Extending these observations to treatment interruption following prolonged ART, we found that CD14^+^ monocytes significantly associated with time to viral rebound at early time points, including AHI, and after 60 weeks of ART treatment. However, their transcriptional profiles became relatively stable at pre-ATI, suggesting a diminished role in late ART-dependent immune variability. In contrast, adaptive immune compartments, especially naive CD8^+^ T cells, emerged as the primary source of transcriptional dynamism during prolonged ART. This shift highlights a transition from early myeloid-driven immune activation toward adaptive immune reprograming over time, positioning naïve CD8^+^ T cells as key regulators of immune readiness and viral rebound potential following treatment interruption.

Although naïve CD8^+^ T cells have traditionally been regarded as a homogeneous quiescent pool, accumulating evidence demonstrates substantial heterogeneity within this compartment^57,58^, as well as progressive transcriptional diversification with age and immune experience^45,59,60^. Consistent with these observations, our study identified a transcriptionally and epigenetically distinct subset within the naïve CD8^+^ T cell pool that emerged along a defined differentiation trajectory during ART. CD8^+^ TNp emerged at AHI and was localized directly adjacent to the interferon-responsive population C13 along the early differentiation trajectory. As ART suppressed viremia, CD8^+^ TNp expanded while interferon signaling diminished. This transition coincided with a marked increase in TNFα-NF-κB and PI3K-AKT-mTOR pathways and AP-1– associated chromatin accessibility, along with their positioning adjacent to T_SL-M_. The enriched AP-1 and TNFα-NF-κB pathways act cooperatively to fine-tune immune activation^61–65^, governing T cell survival, metabolic fitness, and activation thresholds^66–70^.

It is worth noting that prior studies demonstrated that TCF1^+^ CD8^+^ HIV-1–specific CD8^+^ T cells have been linked to durable viral control and long-term immune persistence^29,30,71^. While CD8^+^ T_SL-M c_ells in this study are defined within the bulk CD8^+^ T cell compartment, they represent a memory compartment characterized by high surface protein expression of CD45RO and the activation-associated marker CD44. These cells retain stem-like features, including transcriptomic expression of stemness-associated markers such as *SELL*, *LEF1*, *CCR7*, together with a characteristic co-expression pattern of intermediate expression of *TCF7* (TCF-1) and *PDCD1* (PD-1). These features position T_SL-M_ within a memory-differentiated state that retains elements of stemness. Here we demonstrate that CD8^+^ T_SL-M_ cells are not defined by antigen specificity and can be readily identified within the total CD8^+^ T cell compartment without the need for *ex vivo* antigenic stimulation. This antigen-agnostic approach obviates the requirement for *in vitro* expansion or functional manipulation, thereby enabling robust quantification from limited sample input and enhancing feasibility in clinical settings where cellular material is constrained. We were thus able to detect this population across multiple HIV-1 clinical outcomes and in both acute and chronic infection, indicating stability across disease stages. Importantly, these cells can also be induced by therapeutic vaccination during ART, highlighting their inducibility and potential plasticity. Notably, higher T_SL-M_ levels were also associated with delayed rebound.

Together, these findings support a unified model in which ART promotes a CD8^+^ T cell differentiation continuum from interferon-primed naïve cells, through a poised intermediate state (TNp), to a stem-like memory compartment (T_SL-M_). Within this axis, TNp cells act as a critical transitional population linking early inflammatory cues to the establishment of durable memory potential. This TNp to T_SL-M_ trajectory is also observed in the setting of chronic infection, providing a mechanistic framework for understanding how early immune activation is reprogrammed into a sustained state of immune preparedness that shapes viral rebound dynamics following treatment interruption. During ART, the CD8^+^ TNp population, representing a transcriptionally poised CD8^+^ T cell state, predicted delayed rebound. Conversely, under conditions of enhanced immune activation, such as therapeutic vaccination, protective signatures shift toward the CD8^+^ T_SL-M_ cell compartment, reflecting further stabilization of this primed state into a durable, memory-competent program. Notably, the transition from TNp to T_SL-M_ was marked by enhanced TNFα–NF-κB signaling, positioning these populations as sequential checkpoints of immune readiness and execution. Together, these data establish a unified framework in which the degree of antigenic and inflammatory pressure determines the dominant CD8^+^ T cell state associated with favorable HIV outcomes, positioning this differentiation axis as a scalable and biologically grounded predictor of post-treatment control with potential for therapeutic modification to improve immunologic control of HIV.

## Materials & Methods

### Study design and participants

Demographic and clinical data, including CD4^+^ T cell count, CD8^+^ T cell count, plasma VL, total HIV DNA, Fiebig stage, and time to VLS, were available for 27 donors (RV411, RV397, RV409, and RV405)^6,7,36,37^ who were initially diagnosed at AHI, and subsequently enrolled in the ATI study, without additional interventions, from the RV254 Thai cohort (**Supplementary Table 1**). For discovery analyses, CITE-seq, ASAP-seq, and flow cytometry were performed on initially cryopreserved PBMC, and plasma was collected at AHI and ART timepoints for proteomic profiles. CITE-seq was also performed on PBMC of 16 participants with CHI enrolled in the Africa cohort study (AFRICOS, RV329) at two timepoints (Baseline CHI, and 3 years after ART suppression)^55^. All participants provided informed consent, and use of samples for research was approved by ethical review boards at the Walter Reed Army Institute of Research, USA, Chulalongkorn University Faculty of Medicine, Thailand, and the Kenya Medical Research Institute, Kenya. Previously published single-cell datasets from RV254 and ACTG A5354 (GSE220790 and GSE256089) with HIV DNA measurements were used for validation^18^.

### Plasma proteomics

Plasma samples from AHI and ART timepoints were analyzed using the SOMAscan® Assay v5.0 (SomaLogic), which quantitates ∼11,000 proteins via SOMAmer-based aptamers. Data were normalized using hybridization controls, median signal normalization, and calibration to correct technical variation. Only human protein targets were retained, resulting in 10,788 aptamers. Limits of detection were addressed using a robust estimate^72^, subtracting an estimated limit of detection (eLoD) from each sample, with negative values set to 0. Quality control removed outlier samples and low-signal or low-reproducibility proteins. Statistics analyses were performed through SomaLogic DataDelve™ Statistics. Exploratory data visualization and statistical evaluations were conducted directly within the built-in SomaLogic DataDelve Statistics tool. To evaluate longitudinal proteomic shifts, log-transformed RFU data were analyzed via paired statistical testing. Multiple testing corrections were handled automatically within the software framework to identify significantly altered proteins between the AHI and ART stages.

### Multiparameter flow cytometry

Freshly thawed PBMC from 18 participants from RV254 (RV411, RV397, RV409) were first stained with the LIVE/DEAD Fixable Aqua Dead Cell Stain Kit (Thermo Fisher Scientific L34957) at room temperature for 10 minutes, then cell surface markers with Alexa Fluor 700-labeled anti-CD3 (BD Biosciences, 561027), BUV496-labeled anti-CD4 (BD Biosciences, 569179), BV785-labeled anti-CD8 (BioLegend, 301046), allophycocyanin H7-labeled anti-CD45RA (BD Biosciences, 560674), PE Cy7-labeled anti-CCR7 (BioLegend, 353226), and BUV737 labeled anti-CD27 monoclonal antibodies (BD Biosciences, 569714) at 4 °C for 20 minutes. Following surface marker staining, cells were washed twice with PBS containing 2% fetal bovine serum (washing buffer), and then fixed and permeabilized with Foxp3/ Transcription Factor Staining Buffer Set (Thermo Fisher Scientific, 00-5523-00). Cells were then washed, stained intracellularly, washed again, and analyzed using a LSRII or FACS Symphony A5 (BD Biosciences). Data were analyzed using FlowJo v.10 or higher (BD Biosciences). We defined CD8^+^ T cells based on the available staining antibodies. CD8^+^ TN were CD45RA^+^CCR7^+^CD27^+^, CD8^+^ T_TD_ were CD45RA^+^CCR7^-^CD27^-^, CD8^+^ T_CM_ were CD45RA^-^CCR7^+^CD27^+^, CD8^+^T_TM_ were CD45RA^-^CCR7^-^CD27^+^, and CD8^+^ T_EM_ were CD45RA^-^CCR7^-^CD27^-^(gating strategies are shown in **Supplementary Figure S6**).

### Library Preparation and Sequencing

Participant PBMCs from three longitudinal time points, pre-ART(AHI), ART, and pre-ATI, were washed, resuspended in Cell Staining Buffer (BioLegend), and divided for concurrent processing using single-cell CITE-seq^34,73^ and ASAP-seq^35^ approaches. For CITE-seq, 8-10 uniquely hashed samples were pooled into batches, washed, and stained with Human Universal Antibody Cocktail v1.0 (both TotalSeq-C; BioLegend) per manufacturer’s suggestions. Using the Chromium NextGEM Single Cell 5′ v1.1 kit, sample batches were added to 4 wells of Chromium Chips G, to target recoveries of 16,000 cells per well, and then loaded into a Chromium Controller (all 10x Genomics) for cell partitioning. Both gene expression (GEX) and surface-expressed protein (PROT) libraries were generated as per manufacturer’s instructions. One sample, PID 0221 at the pre-ATI timepoint, was found to have major RBC contamination during preparation and was not sequenced. For ASAP-seq the antibody staining and batching approaches were similar to the above but with TotalSeq-A antibodies (BioLegend). Subsequent cell fixation and permeabilization steps were performed as described previosuly^35^. Using the Chromium NextGEM Single Cell ATAC v1.1 kit, sample batches were added similarly to Chromium Chips H, to target recoveries of 20,000 cells per well (both 10x Genomics). Chromatin accessibility (NUCL) and PROT libraries were generated per manufacturer’s instructions with specific adjustments^35^. Libraries were assessed for quality and concentration with the DNA High Sensitivity kit on the BioAnalyzer (both Agilent), pooled, and quantitated using a MiSeq Nano Reagent Kit v2 and MiSeq instrument (both Illumina). Sample library pools were re-balanced and sequenced using the NovaSeq 6000 S4 Reagent Kit (300 cycles) on a NovaSeq 6000 instrument (both Illumina).

### Single-cell data preprocessing and quality control

Single-cell gene expression data from CITE-seq was generated using the 10x Genomics Cellranger pipeline (v6.1.2) (cellranger count) and the 10x Genomics human reference library (GRCh38 and Ensembl GTF v97)^18,43^. For the hashed sequencing runs, the average number of genes per cell was 1292 and the average number of unique molecular identifiers (UMIs) for RNA transcripts was 3648. The mean read depth per cell was approximately 24,000-51,000 reads for the gene expression library and 9,000-20,000 reads for the antibody library. The minimum fraction of gene expression reads mapped to the genome was 86.4% and RNA sequencing saturation was, on average, greater than 64%.

Single-cell chromatin accessibility data from ASAP-seq was generated with the 10x Genomics Cellranger pipeline (cellranger-arc count) using GRCh38 as reference. Counts for hashed protein libraries were computed. The average read pairs per cell were 48563, while the average percentage of reads mapped to the genome was 76.48%, with a mean sequencing saturation of 61.68%. Downstream analyses of Cellranger outputs were performed using the R package Seurat (v4.3.0); we removed 1 outlier sample (PID 0198 at the ART timepoint), which had < 2000 cells. Finally, we selected only the 2000 most highly variable genes, using the standard procedure in Seurat. HTO expression matrices were normalized using CLR, followed by demultiplexing using Seurat, and then assigned to specific participants using the methods described^43,44^. Negative cells, hash doublets and cells with greater than 10% mitochondrial gene expression were removed. Gene expression matrices (containing a total of 36,602 genes) for all 27 PLWH were log normalized. Antibody -derived tag (ADT) expression matrices for surface protein profiling were incorporated as a separate multimodal assay within the Seurat object and normalized using CLR across cells (margin = 2). A total of 433,525 cells passed the above quality control criteria. ASAP-seq count matrices were processed using Seurat (v4.3.0) and Signac (v1.4.0). Peaks greater than 10000 bp and smaller than 20 bp were filtered out. Single-cell ATAC-seq profiles were filtered to retain high-quality cells based on the following criteria: 3,000–20,000 fragments mapping to peak regions (peak_region_fragments), >15% of reads located within called peaks (pct_reads_in_peaks), a blacklist ratio < 0.05, nucleosome signal < 4, and transcription start site (TSS) enrichment >2. Singlet cells were identified based on HTO barcodes using Seurat’s HTODemux function with default parameters. Surface protein expression from the ASAP-seq dataset was added as a separate assay to the Seurat object and subjected to CLR normalization cross cells. A total of 305,487 cells passed above quality control criteria.

### Integration, dimensionality reduction and annotation

For the CITE-seq dataset, cells were grouped into ‘samples’, defined as ‘participant + timepoint’ combinations. To evaluate confounding batch effects and identify statistical outliers, principal component analysis (PCA) was performed using the averaged gene expression profiles cross individual samples. We then performed reference-based integration, splitting the object by sample and defining integration anchors using the top 2,000 variable features with the first sample as the baseline reference, followed by integrating using the set of anchors. Cells were clustered using the WNN method (resolution: 0.3), using gene & protein features concurrently, and annotated using canonical lineage markers and differentially expressed genes and proteins. For visualization, the dimensionality was reduced using uniform manifold approximation and projection (UMAP) with Seurat in R, selecting the first 30 dimensions. The principal components used to calculate the embedding were the same as those used for clustering. Additional QC was performed to remove cells identified as RBCs (by expression of ‘HBB’), doublets (identified as more than one cell type, or having no clear markers). Furthermore, samples from participants PID 0003, 0114, 0128 and 0243 at AHI timepoint, PID 0198 at the ART, PID 0221 at pre-ATI were excluded due to failure in quality control metrics. After filtering, a total of 128,752 cells were retained at AHI (week 0, baseline), 145,994 cells at ART (60 weeks post baseline); and 137,556 cells at pre-ATI. Cell communication analyses were performed by CellChat package^38^, to examine the communication between major cell types in AHI, ART, and pre-ATI timepoints. Cell-cell communication networks were constructed by merging significant ligand-receptor pairs and their associated degrees.

For the ASAP-seq dataset, cells were clustered using Latent Semantic Indexing (LSI) (resolution 0.8); followed by dimensionality reduction using UMAP with Signac in R, selecting dimensions 2 through 30. Cell annotations were transferred from CITE-seq data using Seurat’s FindTransferAnchors and TransferData; performed using a ’gene activity matrix’ created by calculating counts per cell in gene body and promoter regions, with default parameters; followed by correlation analyses with gene expression levels in the CITE-seq data.

### Differential gene and accessibility analyses

Differential gene expression analyses were performed using the MAST statistical framework in R^40^, modeling gene expression within each cell subset as a function of continuous viral rebound time (days). Genes expressed in fewer than 10% of cells within each subset were excluded to reduce sparsity. Additionally, one participant with HLA-B*57 (PID 0275), who showed no viral rebound during the study, was excluded. The MAST hurdle model was implemented with PID and Time to VL suppression, and statistical significance was assessed for the association between gene expression and rebound time. *P* values were adjusted for multiple testing using the Bonferroni correction. Genes were considered significant if they met both an effect size threshold (|β| > 0.01) and a Bonferroni-adjusted *P* value < 0.05. Genes not meeting these criteria were excluded from downstream analyses. Additionally, a predefined set of genes associated with technical artifacts (e.g., mitochondrial or ribosomal genes) was removed based on a curated blacklist. Differential accessibility was calculated by logistic regression in Seurat. Motif activity was calculated with chromVAR and differential chromatin activity was calculated by comparing mean differences in chromVAR scores using Wilcoxon rank sum tests.

### Trajectory and pseudotime analyses

Trajectory analysis was using Monocle 3^49,50^ was performed on CD8^+^ T cells using UMAP with PCA dimensions 1:20. CD8^+^ T cell subclusters were assigned (resolution: 5e-5), followed by graph-based trajectory learning to construct cell-state lineages. Differential gene expression along pseudotime was assessed using the graph-autocorrelation approach in Monocle 3. Models were fitted for selected gene or protein features along pseudotime for gene and protein features. Curves were predicted from the models and plotted for cells sorted by pseudotime with ComplexHeatmap. To complement the inferences from pseudotime trajectory analyses, PAGA^51^ was applied to the CD8^+^ T cell subset using Scanpy v1.9. Normalized expression matrices and nearest-neighbor graphs were generated, and the connectivity of clusters was computed to model the coarse-grained lineage topology. PAGA graphs were visualized in UMAP space, capturing potential differentiation paths from naïve to memory-like states. Edge confidence values were used to quantify the connectivity between subclusters, supporting the trajectory relationships observed in Monocle 3.

### Gene regulatory network (GRN) analysis

The raw counts from all three timepoints in the CD8^+^ T cell object were used as input for the pySCENIC pipeline.^74^ The pipeline was run using the command line Singularity image version 0.12.1 on the command line. The default grnboost2 method was used to create an adjacency matrix and possible regulators were restricted to transcription factors from allTFs_hg38.txt. Transcription factor regulons based on motif enrichment were found using ranking databases from hg38 v10 (10kbp_up_10kbp_down and 500bp_up_100bp_down), with the HGNC motifs-v9-nr motif to TF annotations database. The regulon activity was calculated from the area under the curve (AUC) regulon enrichment per cell from the counts matrix. The schex package was used to plot the mean AUC per bin (nbins=60, performed separately per timepoint). The AUC for each regulon’s target genes were averaged per subcluster, excluding cluster 15 and 16 due to low cell counts, and converted to z-scores for a selected list of regulons.

### Mapping and annotating query datasets

Independent scRNA-seq datasets from Ehrenberg et al.^18^ and the RV405 and RV329 cohorts, were mapped to the CD8^+^ T cell reference atlas generated in this study. Query scRNA-seq data were processed using Seurat (v5) with standard QC filtering, normalization, scaling, PCA, and CD8^+^ T cell selection. We projected the CD8+ T cell UMAP onto the CD8+ T cell ASAP-seq dataset from the ATI dataset. scATAC-seq data were processed with Signac (v1.12), including filtering by TSS enrichment and nucleosome signal, TF-IDF normalization, and calculation of gene activity scores. Cross-modal integration and label transfer were performed using Seurat’s workflow, projecting query cells into the reference embedding and assigning subcluster identities using TransferData. Mapping accuracy was assessed by prediction scores and marker concordance. Predicted cluster frequencies were quantified per donor and timepoint for downstream comparative and correlation analyses. In addition, two participants from RV405 cohort were excluded from downstream analysis due to atypical clustering and separation in the reference embedding (UMAP) relative to the overall dataset.

### Functional scores and pathway analysis

We used externally-defined lists of genes to determine the functional scores for the CD8^+^ T cells for the following gene sets: naïve^75^, exhaustion^75^, and effector^76^. Module scores were added to each cell using Seurat’s FindModuleScore. The gene lists used can be found in **Supplementary Table 2.**

Gene set enrichment analysis (GSEA) was performed using Hallmark gene sets obtained from MSigDB via the *msigdbr* R package^46^. DEG were filtered to include those with nominal *P* <0.05 and ranked based on direction and *P* value. Pre-ranked GSEA was conducted using the *fgsea* R package with default parameters. For each comparison, normalized enrichment scores (NES) and adjusted *P* value were calculated. Enrichr was used for gene ontology and gene enrichment analysis^77^.

### LASSO regression modeling

Least Absolute Shrinkage and Selection Operator (LASSO) regression was performed to identify the most informative predictors of time to viral rebound.. Since all participants experienced viral rebound during follow-up (i.e., no censoring observation), time to viral rebound was modeled as a continuous outcome, which was log-transformed to reduce skewness. Predictor variables with skewed distributions, including ART duration, pre-ART VL, Time to VLS, pre-ATI CD4^+^ T cell, and pre-ATI CD8^+^ T cell counts, were also log-transformed before modeling fitting. LASSO regression was implemented using the *glmnet* R package, and the optimal penalty parameter (λ) was determined using Leave-One-Out-Cross-validation (LOOCV). The final model was selected using the ‘lambda.1se’ value which identifies the most parsimonious model (i.e., fewest predictors) that is within one standard error of the minimum mean-squared error, thereby shrinking the coefficients of non-impactful predictors to zero.

### Statistical analyses

Correlations were performed by Spearman’s rank correlation coefficient with monotonic lines showing directionality. Comparisons between two groups were performed using nonparametric tests. Specifically, unpaired two-group comparisons were conducted using the Mann–Whitney U test. For matched samples, differences were evaluated using the Wilcoxon matched-pairs signed-rank test. A two-sided *P* value of < 0.05 was considered statistically significant for all statistical analyses. Bonferroni or FDR corrections were applied for multiple testing when appropriate. Survival analyses were performed using the *survival* and *survminer* R package. Optimal cutpoints for outcome-based stratification were determined using the surv_cutpoint function, followed by categorization with surv_categorize. Survival curves were visualized using the *ggsurvfit* package, and group differences were evaluated using the log-rank test implemented via the survdiff function. All descriptive and inferential statistical analyses were performed using R 4.4.3 and higher, and GraphPad Prism 9.0 statistical software packages (GraphPad Software, La Jolla CA). Figures were prepared using GraphPad Prism, R, and Python.

## Data availability

Single-cell multiomics data and code are available in Figshare The GRCh38 reference genome is available from NCBI GenBank (GCA_000001405.15).

## Acknowledgements

We would like to thank the study participants from the RV254/SEARCH 010 cohort, which is supported by cooperative agreements (W81XWH-18-2-0040) between the Henry M. Jackson Foundation for the Advancement of Military Medicine, Inc. (HJF), and the U.S. Department of War (DoW), and in part by the Division of AIDS, National Institute of Allergy and Infectious Diseases, National Institute of Health (DAIDS, NIAID, NIH) (AAI21058-001-01000) and the NIH I4C MDC (5UM1AI64556-05). We would like to acknowledge Dr. Jintanat Ananworanich and Dr. Denise Hsu for their pioneering efforts initiating the RV254 clinical cohort in Thailand. Antiretroviral therapy for RV254/SEARCH 010 participants was supported by the Thai Government Pharmaceutical Organization, Gilead Sciences, Merck and ViiV Healthcare. The views expressed here reflect the results of research conducted by the author(s) and do not necessarily reflect the official policy or position of the Defense Health Agency, Department of War, U.S. Government or HJF. The study protocol was approved by the relevant Institutional Review Board(s) in compliance with all applicable Federal regulations governing the protection of human participants.

## Author contributions

R.T. conceptualized and led the overall project. P.K.E. performed single-cell omics assays, J.W., G.K., A.G., D.E., A.D., T.E., and R.T. contributed to multiomics data analyses and interpretation. H.T. and L.T. performed flow cytometry staining and analysis. C.S., S.S., N.P., S.P., F. S., S.V., N. L.M., and J.A.A provided clinical cohorts and strategy. J.W. and R.T. contributed to the writing of the original manuscript, and all authors reviewed and edited the manuscript.

## References

1 Deeks, S. G., Lewin, S. R. & Havlir, D. V. The end of AIDS: HIV infection as a chronic disease. Lancet 382, 1525–1533 (2013). 10.1016/S0140-6736(13)61809-7

2 Richman, D. D. et al. The challenge of finding a cure for HIV infection. Science 323, 1304–1307 (2009). 10.1126/science.1165706

3 Finzi, D. et al. Latent infection of CD4+ T cells provides a mechanism for lifelong persistence of HIV-1, even in patients on effective combination therapy. Nat Med 5, 512–517 (1999). 10.1038/8394

4. Barouch, D. H. & Deeks, S. G. Immunologic strategies for HIV-1 remission and eradication. Science 345, 169–174 (2014). 10.1126/science.1255512

5 Hocqueloux, L. et al. Long-term immunovirologic control following antiretroviral therapy interruption in patients treated at the time of primary HIV-1 infection. AIDS 24, 1598–1601 (2010).

6 Colby, D. J. et al. Rapid HIV RNA rebound after antiretroviral treatment interruption in persons durably suppressed in Fiebig I acute HIV infection. Nat Med 24, 923–926 (2018). 10.1038/s41591-018-0026-6

7 Colby, D. J. et al. Safety and immunogenicity of Ad26 and MVA vaccines in acutely treated HIV and effect on viral rebound after antiretroviral therapy interruption. Nat Med 26, 498–501 (2020). 10.1038/s41591-020-0774-y

8 Saez-Cirion, A. et al. Post-treatment HIV-1 controllers with a long-term virological remission after the interruption of early initiated antiretroviral therapy ANRS VISCONTI Study. PLoS Pathog 9, e1003211 (2013). 10.1371/journal.ppat.1003211

9 Namazi, G. et al. The Control of HIV After Antiretroviral Medication Pause (CHAMP) Study: Posttreatment Controllers Identified From 14 Clinical Studies. J Infect Dis 218, 1954–1963 (2018). 10.1093/infdis/jiy479

10 Gianella, S. et al. Viral and Immune Risk Factors of HIV Rebound After Interruption of Antiretroviral Therapy. J Infect Dis 231, 1221–1229 (2025). 10.1093/infdis/jiae585

11 Mdluli, T. et al. Acute HIV-1 infection viremia associate with rebound upon treatment interruption. Med 3, 622–635 e623 (2022). 10.1016/j.medj.2022.06.009

12 Li, J. Z. et al. The size of the expressed HIV reservoir predicts timing of viral rebound after treatment interruption. AIDS 30, 343–353 (2016). 10.1097/QAD.0000000000000953

13 Williams, J. P. et al. HIV-1 DNA predicts disease progression and post-treatment virological control. Elife 3, e03821 (2014). 10.7554/eLife.03821

14 Gunst, J. D. et al. Time to HIV viral rebound and frequency of post-treatment control after analytical interruption of antiretroviral therapy: an individual data-based meta-analysis of 24 prospective studies. Nat Commun 16, 906 (2025). 10.1038/s41467-025-56116-1

15 Park, Y. J. et al. Impact of HLA Class I Alleles on Timing of HIV Rebound After Antiretroviral Treatment Interruption. Pathog Immun 2, 431–445 (2017). 10.20411/pai.v2i3.222

16 Prator, C. A. et al. Circulating CD30+CD4+ T Cells Increase Before Human Immunodeficiency Virus Rebound After Analytical Antiretroviral Treatment Interruption. J Infect Dis 221, 1146–1155 (2020). 10.1093/infdis/jiz572

17 Pace, M. et al. Impact of antiretroviral therapy in primary HIV infection on natural killer cell function and the association with viral rebound and HIV DNA following treatment interruption. Front Immunol 13, 878743 (2022). 10.3389/fimmu.2022.878743

18 Ehrenberg, P. K. et al. Single-cell analyses identify monocyte gene expression profiles that influence HIV-1 reservoir size in acutely treated cohorts. Nat Commun 16, 4975 (2025). 10.1038/s41467-025-59833-9

19. 19 Iyer, L. R. & Thomas, R. Current insight into HIV-1 persistence from single-cell transcriptome profiling in acutely treated cohorts of infection. Curr Opin HIV AIDS 20, 481–487 (2025). 10.1097/COH.0000000000000962

20 Aid, M. et al. Follicular CD4 T Helper Cells As a Major HIV Reservoir Compartment: A Molecular Perspective. Front Immunol 9, 895 (2018). 10.3389/fimmu.2018.00895

21 Xu, Y., Ollerton, M. T. & Connick, E. Follicular T-cell subsets in HIV infection: recent advances in pathogenesis research. Curr Opin HIV AIDS 14, 71–76 (2019). 10.1097/COH.0000000000000525

22 Buckner, C. M. et al. Maintenance of HIV-Specific Memory B-Cell Responses in Elite Controllers Despite Low Viral Burdens. J Infect Dis 214, 390–398 (2016). 10.1093/infdis/jiw163

23 Danesh, A., Ren, Y. & Brad Jones, R. Roles of fragment crystallizable-mediated effector functions in broadly neutralizing antibody activity against HIV. Curr Opin HIV AIDS 15, 316–323 (2020). 10.1097/COH.0000000000000644

24 Kaslow, R. A. et al. Influence of combinations of human major histocompatibility complex genes on the course of HIV-1 infection. Nat Med 2, 405–411 (1996). 10.1038/nm0496-405

25 Collins, D. R., Gaiha, G. D. & Walker, B. D. CD8(+) T cells in HIV control, cure and prevention. Nat Rev Immunol 20, 471–482 (2020). 10.1038/s41577-020-0274-9

26 Perdomo-Celis, F., Taborda, N. A. & Rugeles, M. T. CD8(+) T-Cell Response to HIV Infection in the Era of Antiretroviral Therapy. Front Immunol 10, 1896 (2019). 10.3389/fimmu.2019.01896

27 Benito, J. M., Lopez, M. & Soriano, V. The role of CD8+ T-cell response in HIV infection. AIDS Rev 6, 79–88 (2004).

28 Balasubramaniam, M., Pandhare, J. & Dash, C. Immune Control of HIV. J Life Sci (Westlake Village) 1, 4–37 (2019).

29 Kiani, Z. et al. CD8(+) T cell stemness precedes post-intervention control of HIV viraemia. Nature 650, 196–204 (2026). 10.1038/s41586-025-09932-w

30 Peluso, M. J. et al. Correlates of HIV-1 control after combination immunotherapy. Nature 650, 187–195 (2026). 10.1038/s41586-025-09929-5

31 Lim, J. et al. Advances in single-cell omics and multiomics for high-resolution molecular profiling. Exp Mol Med 56, 515–526 (2024). 10.1038/s12276-024-01186-2

32 Tan, W. L. W. et al. Current and future perspectives of single-cell multi-omics technologies in cardiovascular research. Nat Cardiovasc Res 2, 20–34 (2023). 10.1038/s44161-022-00205-7

33 Huang, X. et al. Single-cell multi-omics and machine learning for dissecting stemness in cancer. Brief Bioinform 26 (2025). 10.1093/bib/bbaf566

34 Stoeckius, M. et al. Simultaneous epitope and transcriptome measurement in single cells. Nat Methods 14, 865–868 (2017). 10.1038/nmeth.4380

35 Mimitou, E. P. et al. Scalable, multimodal profiling of chromatin accessibility, gene expression and protein levels in single cells. Nat Biotechnol 39, 1246–1258 (2021). 10.1038/s41587-021-00927-2

36 Crowell, T. A. et al. Safety and efficacy of VRC01 broadly neutralising antibodies in adults with acutely treated HIV (RV397): a phase 2, randomised, double-blind, placebo-controlled trial. Lancet HIV 6, e297–e306 (2019). 10.1016/S2352-3018(19)30053-0

37 Kroon, E. et al. A randomized trial of vorinostat with treatment interruption after initiating antiretroviral therapy during acute HIV-1 infection. J Virus Erad 6, 100004 (2020). 10.1016/j.jve.2020.100004

38 Jin, S. et al. Inference and analysis of cell-cell communication using CellChat. Nat Commun 12, 1088 (2021). 10.1038/s41467-021-21246-9

39 Lundberg, S. M. & Lee, S.-I. in Proceedings of the 31st International Conference on Neural Information Processing Systems 4768–4777 (New York, 2017).

40 Finak, G. et al. MAST: a flexible statistical framework for assessing transcriptional changes and characterizing heterogeneity in single-cell RNA sequencing data. Genome Biol 16, 278 (2015). 10.1186/s13059-015-0844-5

41 Stacey, A. R. et al. Induction of a striking systemic cytokine cascade prior to peak viremia in acute human immunodeficiency virus type 1 infection, in contrast to more modest and delayed responses in acute hepatitis B and & virus infections. J Virol 83, 3719–3733 (2009). 10.1128/JVI.01844-08

42 Iyer, S. S. et al. Resistance to type 1 interferons is a major determinant of HIV-1 transmission fitness. Proc Natl Acad Sci U S A 114, E590–E599 (2017). 10.1073/pnas.1620144114

43 Geretz, A. et al. Single-cell transcriptomics identifies prothymosin alpha restriction of HIV-1 in vivo. Sci Transl Med 15, eadg0873 (2023). 10.1126/scitranslmed.adg0873

44 Li, S. S. et al. HLA-B *46 associates with rapid HIV disease progression in Asian cohorts and prominent differences in NK cell phenotype. Cell Host Microbe 30, 1173–1185 e1178 (2022). 10.1016/j.chom.2022.06.005

45 Gong, Q. et al. Multi-omic profiling reveals age-related immune dynamics in healthy adults. Nature (2025). 10.1038/s41586-025-09686-5

46 Subramanian, A. et al. Gene set enrichment analysis: a knowledge-based approach for interpreting genome-wide expression profiles. Proc Natl Acad Sci U S A 102, 15545–15550 (2005). 10.1073/pnas.0506580102

47 Hao, Y. et al. Integrated analysis of multimodal single-cell data. Cell 184, 3573–3587 e3529 (2021). 10.1016/j.cell.2021.04.048

48 Trapnell, C. et al. The dynamics and regulators of cell fate decisions are revealed by pseudotemporal ordering of single cells. Nat Biotechnol 32, 381–386 (2014). 10.1038/nbt.2859

49 Qiu, X. et al. Reversed graph embedding resolves complex single-cell trajectories. Nat Methods 14, 979–982 (2017). 10.1038/nmeth.4402

50 Cao, J. et al. The single-cell transcriptional landscape of mammalian organogenesis. Nature 566, 496–502 (2019). 10.1038/s41586-019-0969-x

51 Wolf, F. A. et al. PAGA: graph abstraction reconciles clustering with trajectory inference through a topology preserving map of single cells. Genome Biol 20, 59 (2019). 10.1186/s13059-019-1663-x

52 Tibshirani, R. The lasso method for variable selection in the Cox model. Stat Med 16, 385–395 (1997). 10.1002/(sici)1097-0258(19970228)16:4<385::aid-sim380>3.0.co;2-3

53 Li, J. Z. et al. Predictors of HIV rebound differ by timing of antiretroviral therapy initiation. JCI Insight 9 (2024). 10.1172/jci.insight.173864

54 Bosch, R. J. et al. Associations Between Multiple Measures of HIV-1 Persistence in Persons on Suppressive Antiretroviral Therapy. J Infect Dis 225, 2163–2166 (2022). 10.1093/infdis/jiac030

55 Ake, J. A. et al. Noninfectious Comorbidity in the African Cohort Study. Clin Infect Dis 69, 639–647 (2019). 10.1093/cid/ciy981

56 Siliciano, J. D. et al. Long-term follow-up studies confirm the stability of the latent reservoir for HIV-1 in resting CD4+ T cells. Nat Med 9, 727–728 (2003). 10.1038/nm880

57 Marcolino, I. et al. Frequent expression of the natural killer cell receptor KLRG1 in human cord blood T cells: correlation with replicative history. Eur J Immunol 34, 2672–2680 (2004). 10.1002/eji.200425282

58 Azzam, H. S. et al. CD5 expression is developmentally regulated by T cell receptor (TCR) signals and TCR avidity. J Exp Med 188, 2301–2311 (1998). 10.1084/jem.188.12.2301

59 Smith, N. L. et al. Developmental Origin Governs CD8(+) T Cell Fate Decisions during Infection. Cell 174, 117–130 e114 (2018). 10.1016/j.cell.2018.05.029

60 Yu, B. et al. Epigenetic landscapes reveal transcription factors that regulate CD8(+) T cell differentiation. Nat Immunol 18, 573–582 (2017). 10.1038/ni.3706

61 Fujioka, S. et al. NF-kappaB and AP-1 connection: mechanism of NF-kappaB-dependent regulation of AP-1 activity. Mol Cell Biol 24, 7806–7819 (2004). 10.1128/MCB.24.17.7806-7819.2004

62 Jutz, S. et al. Assessment of costimulation and coinhibition in a triple parameter T cell reporter line: Simultaneous measurement of NF-kappaB, NFAT and AP-1. J Immunol Methods 430, 10–20 (2016). 10.1016/j.jim.2016.01.007

63 Fan, H. et al. Oxygen radicals trigger activation of NF-kappaB and AP-1 and upregulation of ICAM-1 in reperfused canine heart. Am J Physiol Heart Circ Physiol 282, H1778–1786 (2002). 10.1152/ajpheart.00796.2000

64 Lee, S. W., Han, S. I., Kim, H. H. & Lee, Z. H. TAK1-dependent activation of AP-1 and c-Jun N-terminal kinase by receptor activator of NF-kappaB. J Biochem Mol Biol 35, 371–376 (2002). 10.5483/bmbrep.2002.35.4.371

65 Huang, W. et al. NFAT and NF-kappaB dynamically co-regulate TCR and CAR signaling responses in human T cells. Cell Rep 42, 112663 (2023). 10.1016/j.celrep.2023.112663

66 Sato, N., Goto, H. & Hirayama, K. [Lactate dehydrogenase isoenzyme in pig dental pulp]. Shoni Shikagaku Zasshi 23, 136–139 (1985).

67 Mehta, A. K., Gracias, D. T. & Croft, M. TNF activity and T cells. Cytokine 101, 14–18 (2018). 10.1016/j.cyto.2016.08.003

68 Waickman, A. T. & Powell, J. D. mTOR, metabolism, and the regulation of T-cell differentiation and function. Immunol Rev 249, 43–58 (2012). 10.1111/j.1600-065X.2012.01152.x

69 Konjar, S. & Veldhoen, M. Dynamic Metabolic State of Tissue Resident CD8 T Cells. Front Immunol 10, 1683 (2019). 10.3389/fimmu.2019.01683

70 Uhl, L. F. K. et al. Interferon-gamma couples CD8(+) T cell avidity and differentiation during infection. Nat Commun 14, 6727 (2023). 10.1038/s41467-023-42455-4

71 Takata, H. et al. Long-term antiretroviral therapy initiated in acute HIV infection prevents residual dysfunction of HIV-specific CD8(+) T cells. EBioMedicine 84, 104253 (2022). 10.1016/j.ebiom.2022.104253

72 Pietzner, M. et al. Synergistic insights into human health from aptamer-and antibody-based proteomic profiling. Nat Commun 12, 6822 (2021). 10.1038/s41467-021-27164-0

73 Stoeckius, M. et al. Cell Hashing with barcoded antibodies enables multiplexing and doublet detection for single cell genomics. Genome Biol 19, 224 (2018). 10.1186/s13059-018-1603-1

74 Van de Sande, B. et al. A scalable SCENIC workflow for single-cell gene regulatory network analysis. Nat Protoc 15, 2247–2276 (2020). 10.1038/s41596-020-0336-2

75 Guo, X. et al. Global characterization of T cells in non-small-cell lung cancer by single-cell sequencing. Nat Med 24, 978–985 (2018). 10.1038/s41591-018-0045-3

76 Herbst, R. S. et al. Predictive correlates of response to the anti-PD-L1 antibody MPDL3280A in cancer patients. Nature 515, 563–567 (2014). 10.1038/nature14011

77 Kuleshov, M. V. et al. Enrichr: a comprehensive gene set enrichment analysis web server 2016 update. Nucleic Acids Res 44, W90–97 (2016). 10.1093/nar/gkw377

